# Using Cancer Profiles to Identify Synthetic Lethal Therapeutic Targets and Predictive Biomarkers in Cancer Gene Dependency Data

**DOI:** 10.1101/2025.04.07.647559

**Authors:** Laurence H. Pearl, Frances M.G. Pearl

## Abstract

Large scale loss-of-function screens utilising CRISPR or siRNA can provide profound insights into the importance of individual genes for the survival of a cancer cell and can drive the identification of therapeutic targets and biomarkers, and the development of targeted drugs. However, the analysis of these data and the substantial bodies of metadata that relate to them, is technically challenging and typically requires substantial expertise in data science and computer coding. To facilitate the analysis of cancer gene dependency data by cancer biologists and clinical scientists, we have developed DepMine – a computational toolkit providing a powerful system for framing complex queries relating cancer gene dependency to the underlying genetic changes that occur in cancer cells. DepMine identifies synthetic lethal relationships between putative target genes and complex ‘cancer profiles’ built from user-specified combinations of mutations, copy-number variation, and expression levels, and can refine these to optimal biomarker definitions for target dependency.

## INTRODUCTION

It is estimated that between 1,000 to 4,000 genes are essential for the survival of a human cell, depending on the method used to define essentiality (Funk *et al*, 2022; Larrimore & Rancati, 2019; Rancati *et al*, 2018). In cancers, as tumours develop and acquire genetic and epigenetic alterations, the tumour cell rewires its regulatory pathways and different sets of genes become essential. Examples of acquired essential genes have been identified by large-scale loss-of-function (LoF) genetic screens (e.g. DRIVE (McDonald *et al*, 2017) and DepMap (Tsherniak *et al*, 2017)).

The DepMap dataset, in which the mutational status and expression profiles of >19,000 genes in >1000 different cell lines has been measured, is the largest amalgamation of data for this. Although not universally true, the dependency of a tumour cell on a gene as determined by CRISPR screens, generally mirrors the susceptibility of that cell to small molecule drugs targeted against the product of that gene (Goncalves *et al*, 2020). Consequently, systematic analysis of the data from such screens is likely to be of great value in development of novel precision therapies and biomarkers.

As cancers evolve, genetic changes are positively selected that directly impact the function of specific genes, with important consequences for the biology of the mutated tumour cell. Typically, these consist of loss-of-function (LoF) mutations in tumour suppressors or gain-of-function (GoF) mutations in proto-oncogenes. LoF of tumour suppressors can be targeted by synthetic sickness/lethality (SSL), a negative genetic interaction whereby a cell can survive the loss of protein product from the tumour suppressor or from its synthetic lethal partner gene, but not from both (Lord & Ashworth, 2017; O’Neil *et al*, 2017). The best-known example is treatment of BRCA1/2 deficient tumours in breast, ovarian and other tumours with PARP inhibitors (Mateo *et al*, 2019). Generally, SSL relates pairs of genes, with one gene being the therapeutic target in cells where the other gene is defective. However, SSL may also relate a target gene to a phenotype or complex genetic state as in the identification of Werner syndrome ATP-dependent helicase (WRN) as a synthetic lethal (SSL) target in tumours displaying microsatellite instability (MSI) (Chan *et al*, 2019; Picco *et al*, 2021); a Werner inhibitor (NCT05838768) is now in Phase 1 clinical development for MSI solid tumours (Ferretti *et al*, 2024).

Oncogene addiction occurs when a tumour cell becomes dependent on the presence of an activated proto-oncogene with a gain-of-function (GoF) mutation (Weinstein & Joe, 2008). Oncogenes can often be targeted directly, but the therapeutic response to even highly specific and potent drugs is often very mixed across patient populations. Synthetic dosage lethality (SDL), where activation of a gene (e.g. the oncogene) is synthetically lethal with LoF of another gene, can be utilised to identify biomarkers where oncogene-directed therapy is most likely to be effective (Megchelenbrink *et al*, 2015).

Given the cost and technical challenge posed by mounting ‘wet’ screens to find SSL relationships, there has been substantial interest in developing analysis pipelines that exploit large cancer cell-line or patient genetic and clinical data collections such as TCGA (Liu *et al*, 2018) and COSMIC (Tate *et al*, 2019)) to identify synthetic lethality relationships, and/or targetable oncogenes and response-predictive biomarkers computationally – for example (Benstead-Hume *et al*, 2019; Das *et al*, 2019; Lee *et al*, 2021; Markowska *et al*, 2023; Pacini *et al*, 2024; Ryan *et al*, 2014; Shao *et al*, 2016a; Srivatsa *et al*, 2022; Wang *et al*, 2022; Wooller *et al*, 2024). To date these methods generally focus on identifying synthetically lethal gene pairs.

Here we describe *DepMine*, a computational toolkit for the analysis of cancer gene dependency data to identify synthetically lethal relationships and the genetic backgrounds in which they might best be exploited. *DepMine* goes substantially beyond pair-wise SSL relationships by allowing specification of complex ‘cancer profiles’ which can be used to probe cancer gene dependency data for synthetic lethality with a target gene of interest within cell lines that match the profile. As well as individual gene descriptors, (i.e. GoF or LoF mutations), individual gene or regional copy number variation (CNV), gene expression levels, and cellular properties (e.g. tissue type, mutational signatures, age, sex), can all be combined in the profile definition using a simple set-theory based algebraic formalism. Profiles can be automatically refined genetic backgrounds in which inhibitors or degraders are most likely to be effective. *DepMine* profiles can then be used as biomarkers to select cancer cell lines or patient samples, in which precision therapies of known drug targets, or specific inhibitors of novel drug targets, are most likely to be of therapeutic benefit.

## MATERIALS AND METHODS

### Data Sources

*DepMine* utilises publicly available datasets centred on DepMap 2023Q2 that can be independently downloaded by the user. The source data files are given in the **Supplementary Data**. Expression thresholds are calculated with an ancillary program *expr_scorer.py*, and mutational signatures for the cell lines documented in DepMap are calculated using *SigProfilerAssignment* https://osf.io/mz79v/wiki/home/ (Diaz-Gay *et al*, 2023) with context_type=96 and genome_build=GRCh38. The input .vcf files required by *SigProfilerAssignment* are generated from the DepMap mutation data by an ancillary program, *MakeVCF.py*.

### Cancer Profiles

At the heart of *DepMine* is a set-theory based system in which membership of a set by a cell line is determined by whether or not that cell line matches a user-defined ‘cancer profile’. *DepMine* cancer profiles are logical statements constructed from primitive functions that determine the truth or otherwise of postulates about the tissue origin of the cell line, sex and/or age of the patient from which it was derived, and the genetic status or expression level of one or more genes within that cell line. A function returning the Universal Set – i.e. all the available cell lines - is also provided. Primitive functions can be combined by the set operations : union **|**, intersection **&** and exclusion **^**, and expressions can be fully bracketed to achieve cancer profiles of considerable complexity (see **RESULTS**).

Operationally, postulate functions are represented by a reserved single character prefix operator followed by its operands. These are summarised in **Table 1**.

**Table 1.**

| Operator | Operand | Resultant set | Example |
| --- | --- | --- | --- |
| + | gene | cells where gene is amplified <sup>1</sup> | **+myc** |
| + | cytogenetic locus | cells where $\geq 1$ gene in locus is amplified <sup>1</sup> | **+17q12.4** |
| - | gene | cells where gene is deleted <sup>1</sup> | **-tp53** |
| - | cytogenetic locus | cells where $\geq 1$ gene in locus is deleted <sup>1</sup> | **-9p21** |
| = | gene | cells where gene is diploid <sup>1</sup> | **=snca** |
| ~ | gene | Cells where gene is involved in a gene fusion | **~alk** |
| > | gene | cells where gene is over-expressed <sup>2</sup> | **>erbb2** |
| < | gene | cells where gene is under-expressed <sup>2</sup> | **<tert** |
| ! | gene | cells where gene has a reading-frame-disruptive loss-of-function mutation | **!brca2** |
| / | gene:mut | cells where gene has defined missense mutation<br>'?' matches any change | **/braf:v600e**<br>**/kras:g12?** |
| / | gene:mut,mut[,mut] | cells where gene has one of several defined missense mutations | **/egfr:l834r,t766m** |
| / | gene+ | Cells where gene has an annotated gain-of-function mutation | **/kras+** |
| / | gene- | Cells where gene has an annotated loss-of-function mutation | **/atm-** |
| \$ | gene | cells where gene has no non-synonymous mutations | **\$rb1** |
| { | Cosmic signature<sup>3</sup> | Cells displaying a defined Cosmic mutational signature | **{sbs7a** |
| { | Cosmic signature aetiology | Cells displaying one or more Cosmic mutational signature attributed to a defined aetiology | **{tobacco** |
| # |  | all cells (Universal Set) |  |
| % | tissue | cells derived from tissue | **%breast** |
| * | sex<br>(Male,Female,Unknown) | cells with sex of patient | **\*fem** |
| ; | age<br>(Adult, Paediatric,Unknown) | cells with age of patient | **;p** |
| @ | Filename | externally sourced CSV file of cell ACH codes | **@CHAN-MSI** |
| 1. Definitions of amplification, deletion and diploid status are based on Mina et al 2020, adapted to the $\log_2(\text{CN ratio} + 1)$ formalism used by DepMap/CCLE.<br>2. Over- and under-expression are defined per gene, based on prior analysis of DepMap expression level values across the full population of cell lines for which data is available (see below).<br>3. As defined in (Diaz-Gay et al., 2023). |  |  |  |

The ***prof*** module provides utilities for showing, renaming and deleting profiles, flattening profiles to primitive functions, and for reading, writing and editing profile collections. Additional modules permit analysis of the tissue distribution of defined profiles - ***type*** - and the frequency of Cosmic mutational signatures (Diaz-Gay *et al*., 2023; Sondka *et al*, 2024) within that cell line selection - ***sig***.

Although developed on the DepMap dataset, the *DepMine* cancer profile system is not restricted to the ≈1100 DepMap data for which dependency data is available, and is readily applicable to any public (e.g. TCGA https://www.cancer.gov/tcga, NCI Genomic Data Commons https://gdc.cancer.gov, Hartwig Medical Database https://www.hartwigmedicalfoundation.nl) or private tumour collection where DNA sequence, copy-number and expression level data are available.

### Defining Expression Thresholds

Unlike copy number variation (CNV), where the baseline ‘dosage’ of almost all genes is the same and common thresholds for deletion and amplification can be defined, expression levels vary widely between different genes, so that no simple global definition of relative over- or under-expression can be applied based on absolute expression levels. Instead, *DepMine* assigns over- or under-expression of any gene in a particular cell line by reference to the observed range of expression values of that gene across all available

Analysis of expression ranges across multiple genes reveals a considerable disparity of ranges both in terms of the absolute expression levels they span, and the distribution of expression values across those ranges. We found that 99.6% of the 19193 genes with DepMap expression data drop into one of three expression range types : a) those in which the gene is expressed in most cells with the expression values (expressed as log_2_(transcripts-per-million+1)) approximate to a normal distribution (∼45%); b) a similar number where the majority of cells do not express the gene and where the values for those that do form an extreme positive skewed distribution (∼43%); and c) a smaller set (∼12%) which are bimodal, being effectively superpositions of the other two distributions (**Figure 1a**).

**Figure 1.**
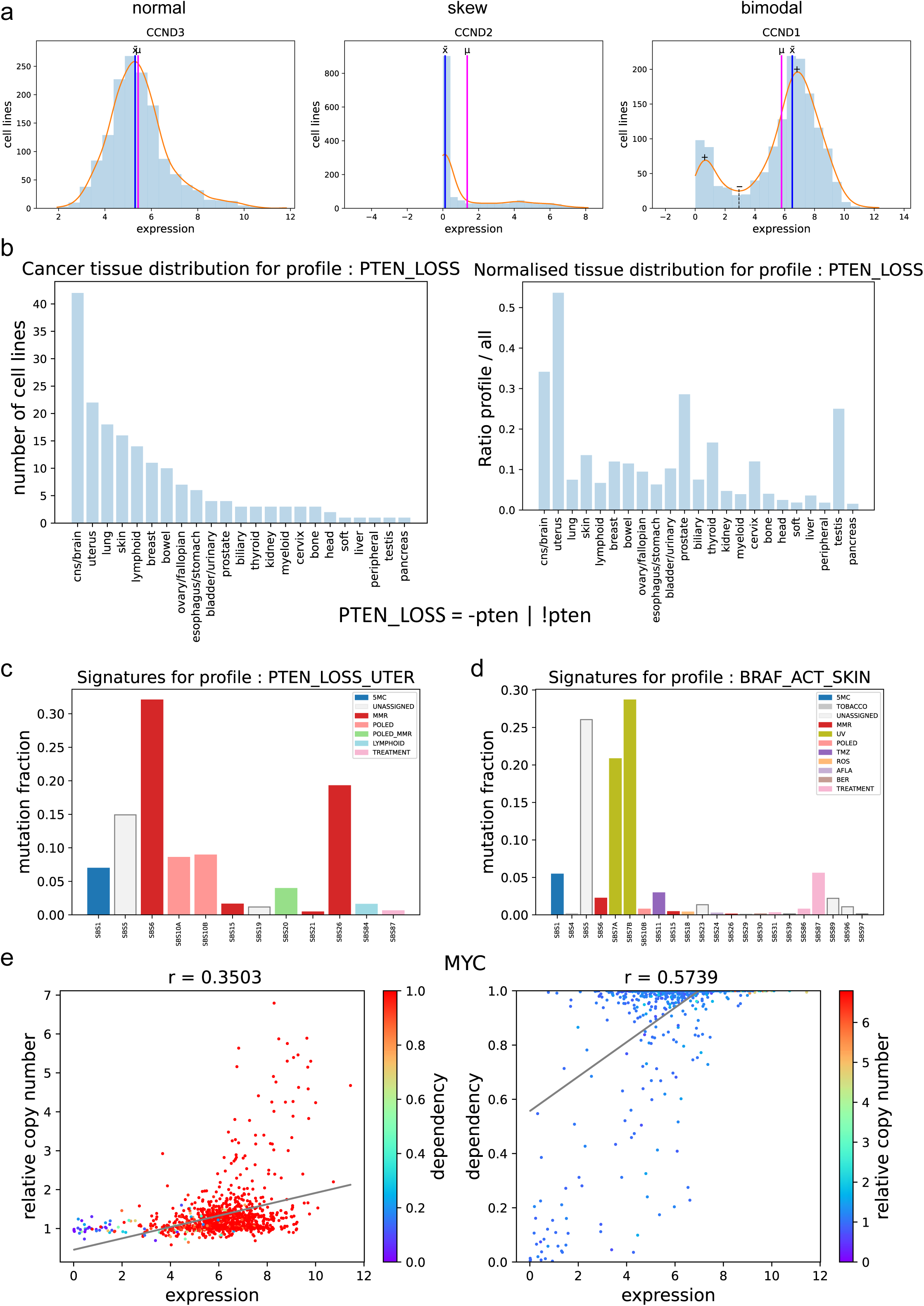
**a)** The three predominant distributions of expression levels −log_2_(transcripts-per-million+1) – in the DepMap data, exemplified by the three paralogues of cyclin D. CCND3 (**leb**) approximates to a normal distribution where the mean (μ) ≈ the median (x°). Numbered lines indicate 1,2 and 3 standard deviations (either side of the mean; CCND2 (**middle**) displays a skew distribution where μ ≠ x° and the modal value is the lowest bin (≈ no expression); and CCND1 (**right**) displays a bimodal distribution in which two maxima (+) and a minimum (−) occur. The orange curve shows the probability density function of the three different distributions calculated using kernel density estimation (KDE). Plots were generated using the **expr** funcRon of DepMine. **b)** Tissue distribuRon of 179 cell lines matching the cancer profile : **PTEN_LOSS** shown (**leb**) as absolute cell line counts, and (**right**) as fracRon of Rssue type normalised by number of that type in the DepMap collecRon. The distribuRon of Rssue types in DepMap is very uneven, so although it appears that PTEN^LoF^ is predominantly in brain tumours, once corrected for the unequal tissue distribution uterine tumours are the most likely to have this geneRc background. The figures are generated by the ***type*** command in the interacRve *DepMine* program. **c)** MutaRonal signature frequencies for cells matching the profile **PTEN_LOSS_UTER** showing strong signals related to mismatch repair deficiency (SBS6, SBS15, SBS26) and/or loss of proof-reading acRvity in replicaRve DNA polymerases (SBS10A, SBS10B, SBS20). **d)** MutaRonal signature frequencies for cells matching the profile **BRAF_ACT_SKIN**, showing a strong signal for mutations resulting from UV-driven DNA damage (SBS7A, SBS7B), as well as SBS5, whose aeRology is unknown. **c**) and **d**) are generated by the ***sig*** command in *DepMine*. **e)** The MYC gene shows a degree of copy number amplificaRon in tumours that is moderately correlated (r=0.35) with expression level (**leb**). High expression of MYC is strongly correlated (0.57) with increased cell dependency (**right**) consistent with its status as an oncogene (see below).

*DepMine* provides useful working definitions for over- and under-expression by assigning bespoke minimum and maximum threshold values for each gene according to the distribution class it belongs to. For bimodal distributions high-expression and low-expression in a given cell line are determined by whether the expression value is respectively greater-than-or-equal to (>=), or less-than-or-equal to (<=) the expression value at the minimum between the two sub-populations, while for skew distributions, high-expression and low-expression are defined as values >= or <= the overall mean of the distribution. For those genes that display an approximately normal distribution, we define the high- and low-expression thresholds as >= μ+2σ and <= μ-2σ, respectively.

Selection of cell lines by expression level uses the primitive postulate functions **>gene** and **<gene** which test against the pre-calculated threshold values for that gene, which in turn depends on which profile type the gene matches (**Figure 1a**). Where a finer degree of control is required, an on-the-fly threshold can be applied by specifying it between the prefix operator and the gene name, e.g. **>3CCND3** will select the very few cell lines in which the CCND3 (cyclin D3) gene is expressed at extremely high levels – >= μ+3σ relative to the rest of the cells in DepMap. More complex expression-based selections can be made using set algebra. e.g.

**CCND3_OUTLIERS = <1ccnd3 | >1ccnd3**

Selects those cells where expression of CCND3 is either > = 1σ above or < = 1σ below the mean expression of CCND1 across the DepMap collection, while :

**CCND3_CENTRAL = # ^ CCND3_OUTLIERS**

uses exclusion from the universal set (generated by **# ^**) to define the effective complement of **CCND3_CENTRAL**, i.e. those cells in which CCND3 is expressed within ± 1σ of the mean.

Graphical interrogation of the expression profile of any specified gene, as in **Figure 1a**, is available via the ***expr*** command in the interactive *DepMine* program

### Identifying Synthetic Sickness/Lethality Relationships

The main goal of *DepMine* is the identification of target genes on which specific tumour cell lines depend for their survival, and of the genetic (and epigenetic) factors in such cell lines that confer dependency on those target genes. Where cellular dependence on one potential target gene (GeneA) depends on the genetic or epigenetic status of a second gene (GeneB), these are deemed to display a Synthetic Sickness/Lethality (SSL) relationship. Identification of simple pairwise SSL relationships for a given GeneA is provided by the ***find*** module of *DepMine*.

To find GeneA-GeneB pairs that show SSL we extract the DepMap gene dependency scores for GeneA across all cell lines, and partition these into one of two classes according to whether GeneB in a given cell line harboured a disruptive (e.g. truncating) mutation and/or a homozygous deletion (CNV < log_2_(0.87/2+1)) and/or showed under-expression (see above for definition) (test class), or whether it harboured no non-synonymous mutations, was diploid (log_2_(2.64/2+1) > CNV > log_2_(1.32/2+1)) and was not under-expressed calculation. These two dependency distributions are then further subdivided according to whether the dependency value is >= or < a threshold dependency (default 0.65, but can be user-specified), and the cell line counts used to populate a 2 x 2 contingency table, which is analysed using a Χ^2^ test (Python **scipy.stats.chi2_contingency**. P-values < 0.05 are taken as indicating a likely SSL. SSL pairs are presented graphically ranked on *p*-value along with the fraction of cell lines in which the GeneB was disrupted/deleted/downregulated, and the magnitude of the effect calculated using Cohen’s d value (Cohen, 1988). As the ***find*** command for a single GeneA can involve more than 20 million statistical computations, it is distributed across all available CPUs using the Python multiprocessing **Pool.starmap** method, so that its execution time depends on the number of CPUs available. For example, where synthetic lethal partners with disruptive mutations were sought for the ALDOC (aldolase C) gene, the ***find*** command took 386 seconds on a 2020 iMac with a 3.8GHz 8-Core Intel Core i7 running OSX 12.6. Results of ***find*** searches are displayed graphically as the kernel density estimates (KDE) of the test and control sets of cells, with distributions means indicated.

Using a highly parallel version of *DepMine* using the 2022Q2 release of DepMap, and running on a multi-node HPC cluster, we performed a complete all-by-all equivalent of the ***find*** command looking for SSL partners with Loss-of-Function (LoF - disruptive mutation *or* CNV deletion) for all of the 16825 genes for which dependency data was available, and which were not genes with high average dependency and therefore unlikely to have utility as therapeutic targets. The results of the all-by-all analysis is discussed below.

*DepMine* extends the definition of SSL relationships through the use of the powerful Cancer Profile system described above. In this mode the ***switch*** module **of** *DepMine* applies the same statistical analysis as the pairwise ***find*** search, but instead looks for GeneAs whose dependency are significantly different in a population of cell lines that match a defined cancer profile as compared with a control set of cell lines. Control sets are generated by selecting cells that have no non-synonymous mutations and are diploid for all the genes specified in the test profile, and have the opposite expression pattern to any genes whose expression is specified in the test profile.

### Mining for Secondary Effects

While analysis of dependency switching using cancer profiles allows identification of target genes with a strong and significant SSL relationship to a complex cancer profile, the compilation of the cancer profiles themselves relies on prior knowledge of the genes whose mutation and/or copy number variation and/or under/over-expression defines the state or phenotype of interest. Thus, in almost all significant and strong SSL relationships identified, as well as a population that matches the test profile and has a high dependency, there will be cell lines that match the test profile, but nonetheless display low dependency on the SSL target. In many ways this mirrors the common observation in cancer clinical trials that there are often a group of patients selected on the same criteria as those who respond to the new therapy, but nonetheless fail to respond. In both cases this is likely to be due to the inadequacy of the profile used to select the cell lines or patients, and begs the question as to how the initial profile, which proves so useful in initially identifying the SSL target or treated patient cohort, can be refined to give a strong predictive ‘biomarker’ for those tumour cell lines (and patients) which are mostly likely to respond to a therapy directed at the identified SSL target. To address this, *DepMine* incorporates a ‘mining’ feature that analyses cell line populations that match the profile, to find secondary features that improve the size of the SSL effect and increase the proportion of the cell lines in the matching distribution showing high dependency on the SSL target.

Secondary mining is implemented in the *DepMine **mine*** module, and allows for analysis of the profile-matching-population on the basis of all the factors that can be used to define cancer profiles : sex, age, tissue type, LoF mutations, GoF mutations, gene fusion, mutational signatures, deletion, amplification, downregulation of expression, upregulation of expression. The ***mine*** algorithm looks for distributions of profile-matching cell lines that additionally match the different secondary factors being tested and determines whether those cells are statistically over-represented in the part of the overall cell line population with low dependency on the SSL target, or are over-represented in the population with high dependency on factor will be expected to improve the size of the SSL effect, while for the latter case, specific inclusion of matching cells will be expected to achieve this.

Inevitably there is a complex interplay and interdependence between the influence different secondary factors have on the effect size, and the effects themselves are not simply additive. So, different combinations of secondary effects need to be explored to find the combination that maximises the predicted effect size, and this may be open to further improvement by a further round (or rounds) of secondary mining. However, a balance needs to be struck between the effect size a cancer profile can achieve and the number of cell lines it ultimately matches. Taken to absurdity, one can envisage an ultimately refined profile that only matches a single cell line, albeit one that has maximal (1.0) dependence on the target gene.

To help find solutions that balance effect size and cell line coverage, *DepMine* provides a Pareto optimisation function (Hu *et al*, 2013) that evaluates a ‘forest’ of all possible decision tree combinations of factors that individually show a strong effect when added to the ‘root’ profile, and identifies those that give an optimal combination of effect size and the number of cell lines they match.

### Cell Line Selection with Cancer Profiles

*DepMine* cancer profiles provide a powerful tool for selecting cell lines on the basis of complex combinations of tissue, age and sex origin of the tumour and the presence of defined mutational signatures, as well as mutational status of genes and genomic loci, and relative expression levels of individual genes. The basic ‘currency’ of SSL analysis - loss of a functional gene product (e.g. PTEN) through deletion or frame-disrupting mutation - can be simply represented and used to analyse the tissue distribution of the loss of that important tumour suppressor (**Figure 1b**). The profile can be further defined to reference only those tumours of uterine origin in which PTEN is lost, and the distribution of mutational signatures in that subset analysed (**Figure 1c**). Activation of an oncogene such as BRAF through gain of function (GoF) mutation is also effectively represented as a *DepMine* profile and combined with a tissue origin, reveals the distinctive UV signature of BRAF-driven melanomas (**Figure 1d**).

Very specific GoF profiles can be developed to encapsulate activation of pathways that can arise due to a variety of genetic events e.g.

**MAPK_ACTIVATED = /hras:g12?,g13?,q61? | /kras:g12?,g13?,q61? | /nras:g12?,g13?,q61? | /raf1:s257l,s259f | /braf+**

defines the set of cells in which any of the RAS family genes have one of the well-characterised oncogenic mutations, or CRAF has either of the missense mutations that preventing its downregulation by 14-3-3 proteins, or has any annotated gain-of-function mutation in BRAF, any or all of which result in an increase in signalling through the canonical RAF-MEK-ERK MAP-kinase cascade.

Selection based on copy number variation e.g. the MYC amplification seen in many tumours,can also be effectively represented, optionally combined with selection based on expression level - e.g. **MYC_UP = +myc | >myc -** although these may often be correlated and redundant. *DepMine* provides a tool in the ***expr*** module to visualise any correlations between copy number and expression, and relates these to the pattern of tumour cell dependency on that gene (**Figure 1e**).

## RESULTS

### Identifying All-By-All SSL Gene Pairs

Using a highly parallel version of DepMine using the 2022Q2 release of DepMap, and running on a multi-node HPC cluster, we performed a complete all-by-all equivalent of the ***find*** command looking for SSL partners with severe LoF (frame disruptive mutation) *or* CNV deletion for all of the 16825 genes for which dependency data was available, and which were not genes with high average dependency and therefore unlikely to have utility as therapeutic targets. We found 34256 GeneA:GeneB pairs where the |dep(GeneA)| < 0.65, and dep(GeneA) in cells with LoF/CNV-del of GeneB was significantly > than GeneB-WT cells (p < 0.05). A full-listing of the SSL pairs identified is provided in **Supplementary Database 1.**

A number of paralogous pairs gave strong and highly significant signals in the pairwise all-by-all LoF analysis, several of which have been previously noted in experimental screens (Ryan *et al*, 2023b). For example, cells in which the SWI/SNF chromatin complex component ARID1A was deleted or had a LoF mutation, showed a significantly stronger dependence on its paralogue ARID1B, which is generally dispensable in cell lines with functional ARID1A (**Figure 2a**). A similar effect was seen with the SWI/SNF helicase SMARCA2 which displays strong dependency in cells with LoF in the paralogous SMARCA4 (**Figure 2b**). These SSL relationships are not generally commutative, so that while loss of ARID1A or SMARCA4 in tumour cells enhances dependency on ARID1B or SMARCA2 respectively, loss of ARID1B or SMARCA2 had little effect on the requirement for ARID1A or SMARCA4 respectively for tumour cell viability (**Figure 2a,b**). However, whether this reflects underlying biology or is the result of small numbers of observed mutations for the second gene in the cell line data, is unclear. The homologous lysine acetyltransferase genes CREBBP and EP300 also display a paralogue SSL relationship which again shows a strong polarity of effect, with tumour cell LoF of EP300 required rather than CREBBP (**Figure 2c**). It is also worthy of note, that even in the orientations that display the effect, these paralogue SSL relationships are not absolute and there remain many cell lines lacking the sensitising paralogue that aren’t dependent on the compensating paralogue. Indeed, searching DepMap reveals more than 20 viable cell lines in which functional ARID1A and functional ARID1B are both lost. Clearly there are secondary factors that define the settings in which these paralogue SSLs operate, but these are not revealed by simple pairwise LoF analysis, experimental or computational.

**Figure 2.**
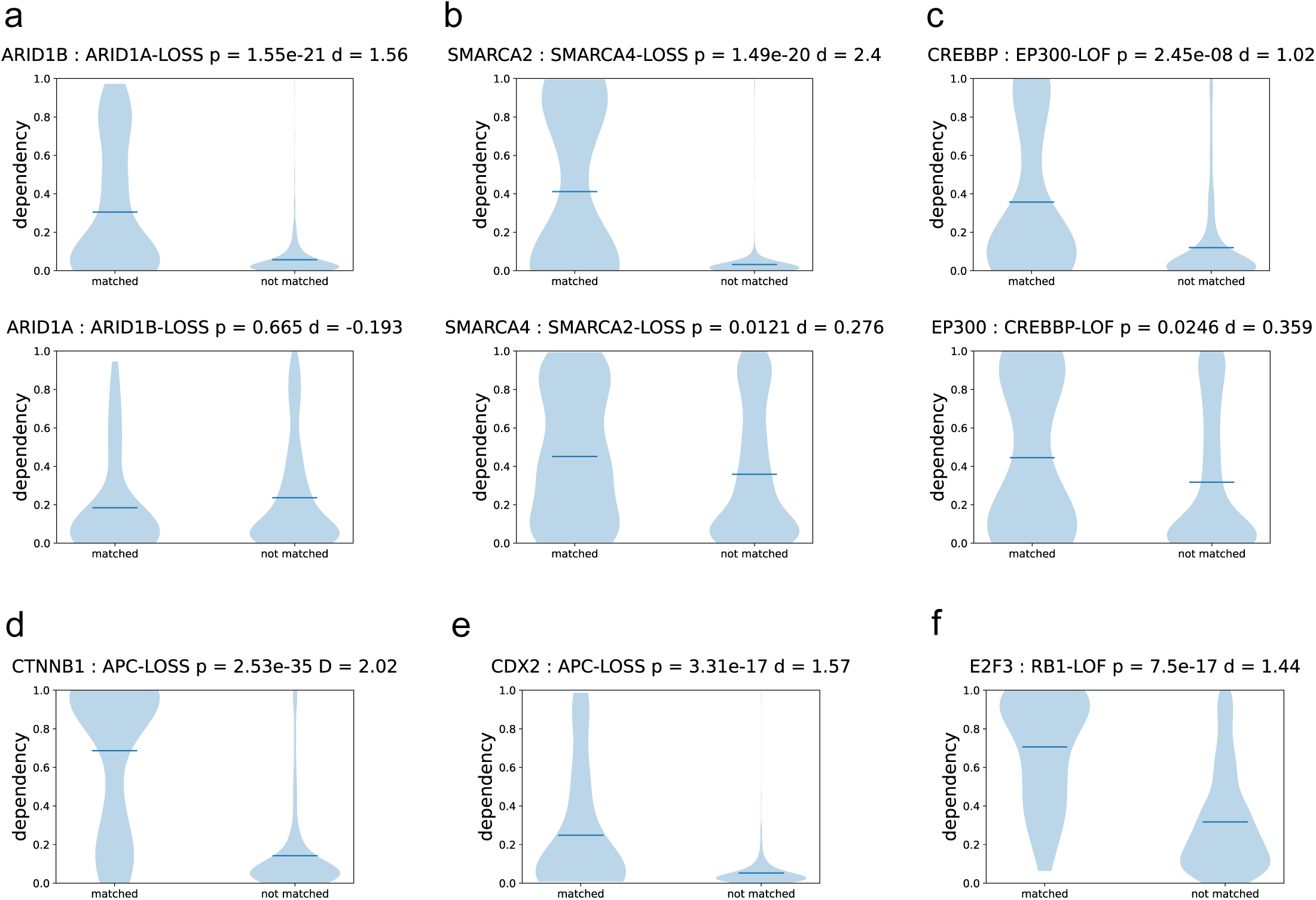
**a)** Kernal Density Estimate (KDE) plots for distribution of dependency on (**top**) ARID1B in cells with ARID1A^LOF^ (matched) compared to cells with ARID1A^WT^ (not matched) – the dark line indicates the mean values of those distributions. The difference is highly significant with a very strong effect (d > 0.8); (**bokom**) as (**top**) but dependency on ARID1A in cells where ARID1B is either LoF or wild-type – unlike (**top**) the distributions show no significant difference. **b)** as **a)** but for SMARCA2 and SMARCA4. **c)** as **a)** but for CREBBP and EP300. **d)** as **a)** but for CTNNB1 dependency in cells with APC^LoF^ compared to cells with APC^WT^. **e)** as **d)** but for CDX2 dependency. **_f)_** as **a)** but for dependency on E2F3 in cells with RB1^LOF^ compared to cells with RB1^WT^

Strong/medium non-paralogous SSL relationships with important tumour suppressors were also identified by the all-by-all pairwise analysis, several of which have already been noted. The most statistically significant SSL pair identified was between CTNNB1 (encoding β-catenin) and deletion/LoF of the adenomatous polyposis cancer associated APC, which is required for regulating β-catenin levels in the Wnt-signalling pathway (**Figure 2d**). APC LoF occurs in ∼30% of sporadic colorectal cancers and correlates with poor prognosis (Li *et al*, 2023). While drugs that aim to block β-catenin’s interactions with downstream factors such as TCF4, BCL9 or CBP have yet to prove useful clinically, the very strong dependence of APC^LoF^ tumour cells on the presence of a functional β-catenin molecule compared to the very low dependence of cells with APC^WT^, suggests that therapeutic depletion of β-catenin using a PROTAC agent could be an effective option with low general background toxicity (Yang *et al*, 2022). Other genes showing medium/strong SSL with APC include CDH1, JUP, TCF7L2 and CDX2 (**Figure 2e**), but none of these offer readily tractable drug targets.

SSLs were identified for LoF of RB1, a common occurrence in cancers with a poor prognosis. The strongest effect was seen with the transcription factor E2F3, whose inhibitory interaction with the RB1 protein regulates entry to S-phase (Chen *et al*, 2009), suggesting that RB1-defective tumours specifically develop a level of ‘oncogene addiction’ to E2F3, but not to other RB1-interacting E2F paralogues which display no significant SSL (**Figure 2f**). The E3-ubiquitin ligase subunit SKP2 also shows a strong SSL with RB1-loss, which has previously been noted experimentally (Gupta *et al*, 2022).

### Unravelling passenger effects in regional deletions

Within the all-by-all pairwise analysis using either frame-disrupting mutations or CNV deletion of GeneBs, a Promiscuous SSL partners included VPS4A, RAB6, HBS1L, KRAS, ITGB1, HSPA13 and WRN. With the exception of WRN (but see later) the multiple GeneBs whose LoF/deletion switched the dependency of these promiscuous GeneAs, showed strong clustering into a small number of chromosomal loci, commonly involving one or both of 9p21 and 18q21 (**Figure 3a**).

**Figure 3.**
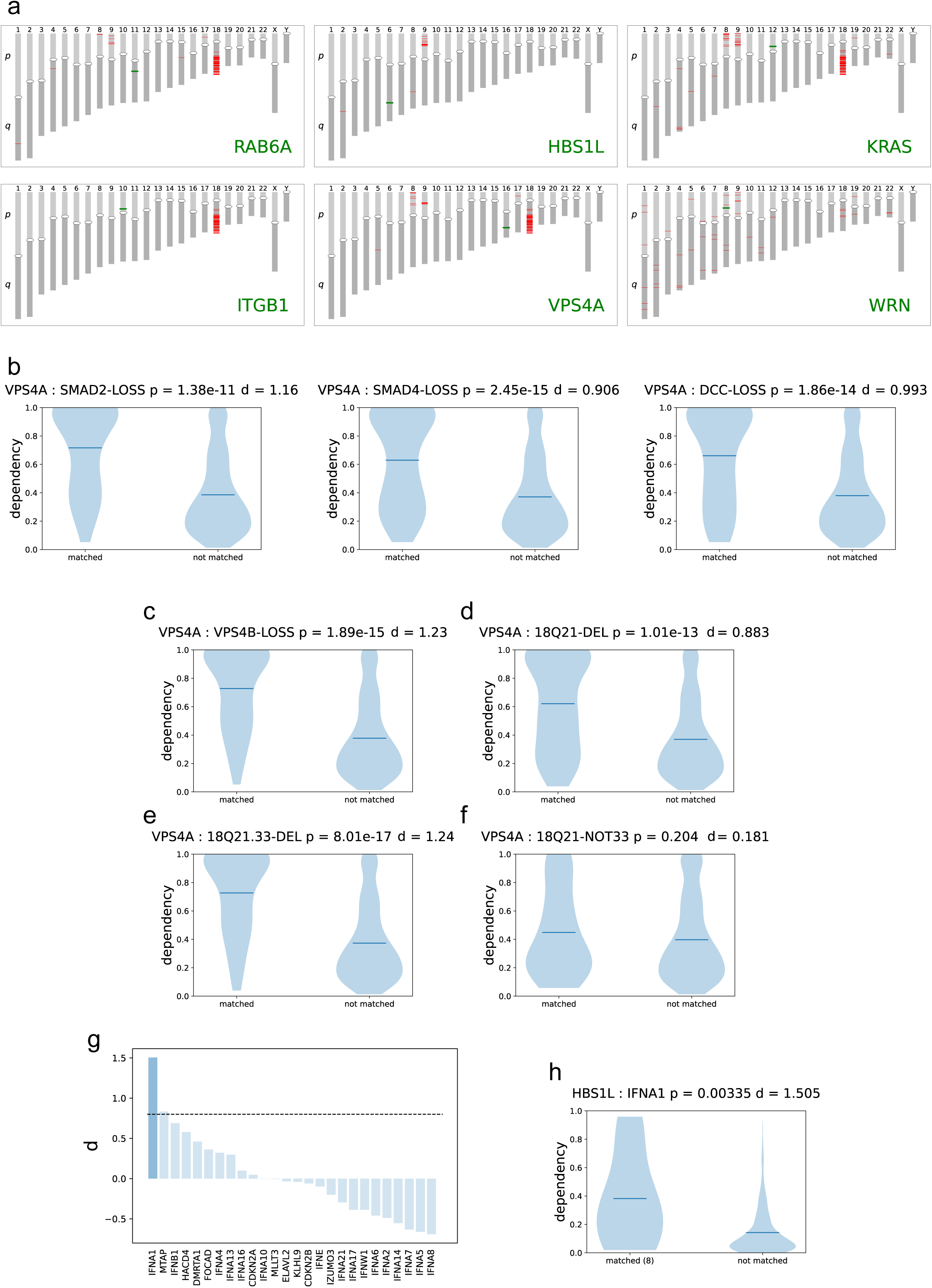
**a)** Chromosomal locaRons (red lines) of the genes whose LoF/deleRon showed a staRsRcally significant SSL effect with one of the promiscuous SSL GeneA partners shown. The locaRon of the GeneA itself is indicated by the thick green line. The positions of centromeres are indicated by white ovals. This plot is generated by the ***chro*** module of *DepMine*. **b)** KDE plots for VPS4A dependency in cells with LoF of (**leb**) SMAD2, (**centre**) SMAD4, or (**right**) DCC compared to cells in which these genes are diploid and harbour no non-synonymous mutations. All three genes show strong, statistically significant SSL effects. **c)** as **b)** but for VPS4A dependency in cells with LoF of VPS4B. **d)** as **b)** but for deletion of any gene(s) in cytogenetic band 18q21. **e)** as **d)** but for deletion of any gene(s)in sub-band 18q21.33. **f)** as **d**) but for deletion of any gene(s) in band 18q21 apart from sub-band 18q21.33 **g)** Waterfall plot of d values for switching of HBS1L dependent on only LoF (missense or frameshit) mutations, but not deletion, of genes in the 9p21.3 cytogenetic locus for which mutations are recorded in DepMap. Dark bars indicate statistical significance (p < 0.05) Only LoF of IFNA1 produces a significant effect > 0.8 – the usual threshold for a strong signal. The plot was generated by the **locu** module in DepMine **h)** KDE plot of HBS1L dependency in cell line with IFNA1 LoF but not deletion, compared with cell lines in which IFNA1 has no mutaRon or deleRon

The power of *DepMine*’s cancer profiles to unravel complex synthetic lethal relationships is substantial. For example : strong SSL relationships were identified in the all-by-all pairwise analysis (see above and **Supplementary Database 1**) between VPS4A and LoF/deletion of a number of other genes, including important tumour suppressors such as SMAD2, SMAD4 and DCC which are disrupted in a range of cancers (**Figure 3b**). While no obvious function or role links the VPS4A gene product, a AAA+-ATPase involved in vacuolar sorting, with the various genes with which it displays an SSL effect, those genes share a common chromosomal location, with the majority mapping to cytogenetic band 18q21 on the long arm of chromosome 18 (**Figure 3a**), a genetic locus often deleted in gastrointestinal and pancreatic cancers, and associated with poor prognosis and liver metastases (Tanaka *et al*, 2006).

At first sight this appears to be a promising ‘hit’ as VPS4A is a druggable target which could be used to treat cancers where SMAD2/SMAD4 signalling has been lost. However, one of the other genes in the 18q21 region is VPS4B, a VPS4A paralogue, with which it has a well-documented SSL relationship (Neggers *et al*, 2020) readily detected by *DepMine* (**Figure 3c**). It therefore becomes difficult to determine whether the effects seen for SMAD2, SMAD4, DCC and other genes in 18q21 are direct effects due to a genuine functional synthetic lethality with VPS4A, or whether it is because of their proximity to the VPS4B gene which is often lost with them as a ‘passenger’ in cancer-associated deletions of 18q21, and so indirectly reflect the paralogue SSL between VPS4A and VPS4B. Certainly, VPS4A generates a statistically significant and reasonably strong SSL effect when tested against cells with a cancer profile defined by deletion of any gene within 18q21 (**18Q21_DEL**), compared to cell lines without any such deletions (**Figure 3d**). So, a paralogue passenger proximity effect could be operating even though SMAD2, SMAD4 and DCC map to sub-bands 18q21.1 and 18q21.2 at the centromere-proximal end of 18q21, while VPS4B maps to sub-band 18q21.33 at the centromere-distal end of 18q21. This type of conundrum in *DepMine* can be resolved by defining a complex cancer profile **18Q21.33_DEL** that selects cell lines with any genes deleted in 18q21.33, and profile **18Q21_NOT33** that selects cell lines with deletion of genes anywhere in 18q21 except 18q21.33. Consistent with the apparent VPS4A SSL with genes other than VPS4B being primarily due to their collateral loss along with VPS4B in regional deletions, VPS4A shows a significant and strong SSL effect with **18Q21.33_DEL** (**Figure 3e**) comparable to the that with LoF of VPS4B (**Figure 3c**), whereas the SSL effect is completely lost when sub-band 18q21.33 deletions are excluded (**Figure 3f**), even though loss of the SMAD2, SMAD4 and DCC genes would still be present in the matched cell line set. We can therefore conclude that there is unlikely to be any direct functionally related SSL between VPS4A and these important tumour suppressors, but that their chromosomal proximity to the VPS4A paralogue VPS4B nonetheless opens up a potential therapeutic vulnerability in the many tumours in which SMAD2, SMAD4 and DCC loss has been positively selected, so long as the deletion that eliminates these tumour suppressors encompasses 18q21.33 and removes VPS4B as a passenger.

HBS1L – which encodes a GTPase involved in release of stalled ribosomes (Shao *et al*, 2016b) – shows significant SSL relationships with a cluster of genes mapping to chromosome band 9p21.3 – a locus that is lost in a range of tumours (Jiang *et al*, 2023; Spiliopoulou *et al*, 2022) and increasingly associated with resistance to immune checkpoint therapy (Han *et al*, 2021). This locus contains CDKN2A - which encodes the well characterised tumour suppressors p16^INK4A^ and p14^ARF^, MTAP – encoding methyladenosine phosphorylase whose loss is associated with poor survival in glioblastoma and other tumours (Patro *et al*, 2022), and a cluster of 13 homologous genes encoding α-interferon isoforms. As HBS1L has no essential paralogues within this region, it is not obvious which gene or genes lost in the 9p21.3 deletion are responsible for switching HBS1L dependence. To address this, the ***locu*** module of *DepMine* performs a scan of all GeneBs in the locus and calculate the switch effect size on the target gene – HBS1L in this case – based only on mutational LoF (nonsense or frameshift), but not copy number variation (**Figure 3g**). For HBS1L, this scan identifies the IFNA1 gene, encoding interferon-α1, as having a very strong and significant most important in sensitising tumours with this deletion to HBS1L loss, and not the more ‘usual suspects’ CDKN2A or MTAP.

Interestingly, experimental dissection of the 9p21.3 locus has shown that specific loss of the interferon cluster in substantial 9p21.3 deletions in a mouse pancreatic cancer model, plays an important role in facilitating immune evasion and immunotherapy resistance (Barriga *et al*, 2022). The SSL relationship we observe here suggests that HBS1L may be a useful target for that context. In similar analyses (see **Supplementary Data**) KRAS, RAB6A and ITGB1 all show significant SSL with mutational LoF of SMAD4 in the 18q21 locus, but only KRAS shows a ‘strong’ effect.

### Analysis with Cancer Profiles

The use of cancer profiles in *DepMine* extends the use of DepMap data for identifying SSL relationships beyond gene pairs, allowing the identification of genes whose dependency is switched on the basis of sets of factors that define complex states. A good example of this is the phenotype of microsatellite instability (MSI) which manifests as expansion/contraction of short repeating DNA mononucleotide and dinucleotide sequences (Ellegren, 2004) and is particularly associated with highly mutated colorectal, gastric and endometrial tumours (Riedinger *et al*, 2024; Sato *et al*, 2023) as well as some haematological malignancies (Maletzki *et al*, 2013). MSI results from LoF or transcriptional downregulation of the genes encoding components of the DNA mismatch repair system (MMR) (Li, 2008) and/or from loss or defects in proof-reading replicative DNA polymerases (Haradhvala *et al*, 2018). Based on this we constructed a *DepMine* cancer profile for microsatellite instability, **MSI_FULL_DOWN**, which matches 142 cell lines in DepMap across a range of tissue types (**Figure 4a**), but dominated by tumours of lymphoid, bowel and uterine origin.

**Figure 4.**
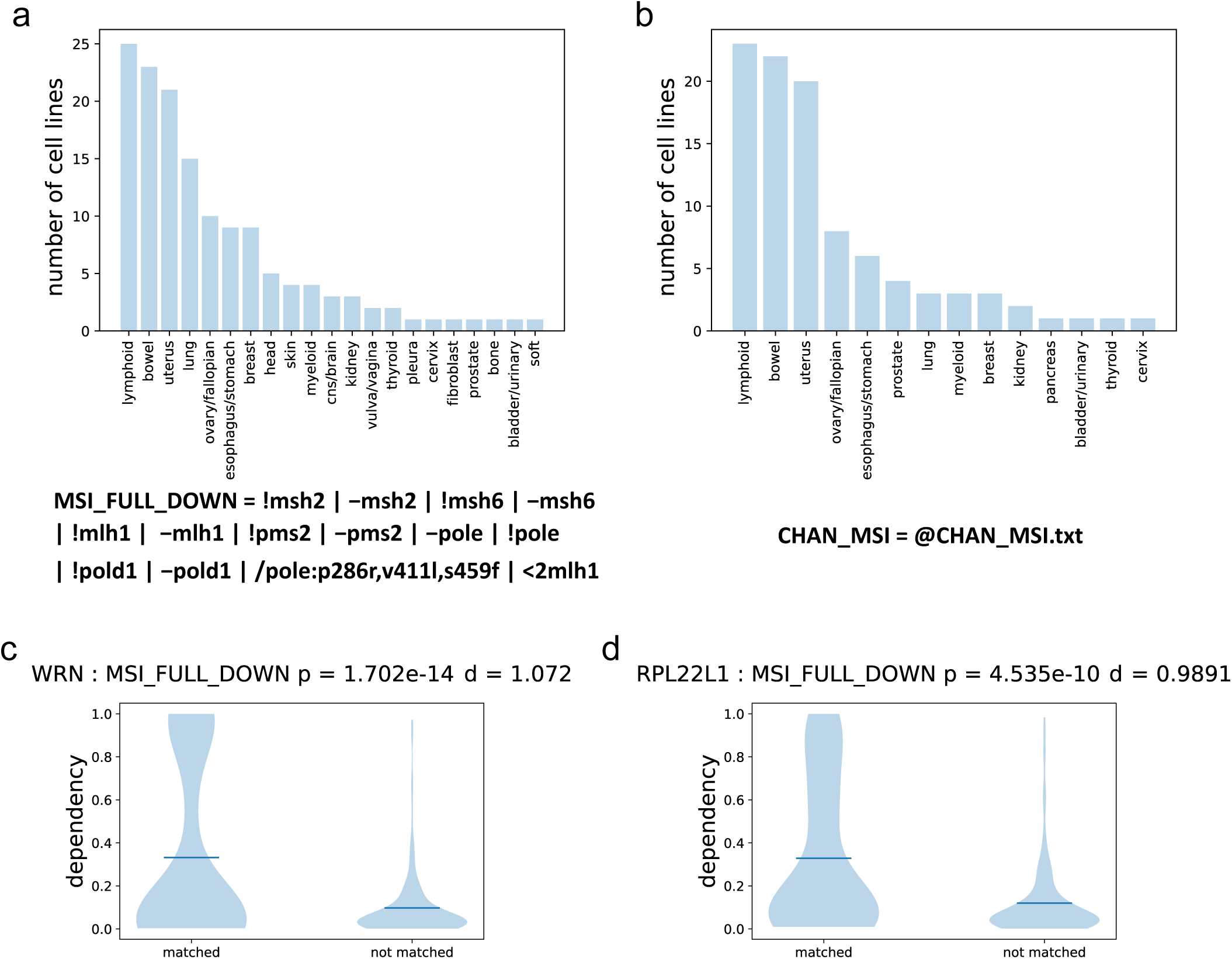
**a)** Tissue distribuRon of 142 cell lines matching the cancer profile : **MSI_FULL_DOWN**. **c)** KDE plot for WRN dependency in cells matching the **MSI_FULL_DOWN** profile compared with a control set of cells **d)** as **c)** but for RPL22L1.

To ‘sense check’ our MSI profile, we defined a second profile (**CHAN_MSI** = **@CHAN_MSI.txt)** by uploading a curated list of cell lines classified as MSI-High based on experimental analysis (Chan *et al*., 2019). **CHAN_MSI** matches 99 cell lines in DepMap also dominated by tumours of lymphoid, bowel and uterine origin (**Figure 4b**), but with a lower representation of lung samples than our computationally defined **MSI_FULL_DOWN** profile. The two profiles have 71 cell lines in common (determined by **CHAN_MSI & MSI_FULL_DOWN**).

There has been considerable interest in identifying druggable targets specific to MSI cells, and a number of experimental CRISPR screens have been performed for this purpose (Chan *et al*., 2019; Picco *et al*., 2021). To determine whether this question was amenable to our computational approaches we utilised the ***switch*** module in *DepMine* which applies a similar analysis to the pairwise ***find*** search, but looks for GeneAs whose dependency is significantly different in a population of cell lines that match a defined cancer profile – e.g. **MSI_FULL_DOWN** as compared with a control set of cell lines. Control sets are generated by selecting cells that have no non-synonymous mutations and are diploid for all the genes specified in the test profile, and have the opposite expression pattern to any genes whose expression is specified in the test profile. Dependency switching analysis using the **MSI_FULL_DOWN** profile found 516 genes displaying SSL with p < 0.05 (-log_10_(p) > 2) and took approximately 9 minutes on a 2020 iMac with a 3.8GHz 8-Core Intel Core i7 running OSX 12.6.

Although many genes displayed statistically significant differences between the dependency distributions for the profile matching and control sets of cell lines, only the highest ranking (based on p-value) two genes also showed strong effects (Cohen d > 0.8). The first of these was WRN – the gene encoding the RECQ-family DNA helicase whose germline mutation underlies the cancer-prone progeria Werner’s syndrome (**Figure 4c**) (Monnat, 2010). WRN is the prime target identified in the several laborious experimental screens that have been conducted (Chan *et al*., 2019; Lieb *et al*, 2019; Picco *et al*., 2021; van Wietmarschen *et al*, 2020), so its unambiguous *ab initio* identification here as a strong SSL partner for MSI by a completely hypothesis-free computational analysis in under 10 minutes on a standard desktop computer, completely vindicates our approach. The second gene showing significance and a strong effect was RPL22L1 (**Figure 4d**) which encodes a ribosomal protein that has previously been implicated in progression and treatment resistance in a range of different cancer types (Chen *et al*, 2023; Rao *et al*, 2019; Yi *et al*, 2023), although the mechanism is not understood. WRN and RPL22L1 were also the top two hits when the ***switch*** analysis was repeated with the **CHAN_MSI** profile (results not shown).

### Gain of Function Dependency Switching

While loss of function due to disruption of the reading frame is straightforward to predict based on the DNA sequence of a gene, it is difficult to predict the consequences of missense mutations. While the majority of these will have no significant effect, a minority will cause loss (LoF) or gain of function (GoF) which may have major consequences.

*DepMine* incorporates a dedicated ***GOF*** module that applies the ***switch*** analysis described above, to a curated list of GoF mutations. This can come from DepMap itself, which annotates 197 GoF mutations across 56 different genes (2023Q2 release), or from an extensive GoF dataset we have hand-compiled over several years from literature reports, which contains 1153 mutations across 67 different proteins (based on (Baeissa *et al*, 2017)).

The **GOF** module can operate at the level of the individual mutation, so that independent ***switch*** analyses are run, for example, for KRAS:G12V (**/kras:g12v**), KRAS:G12D (**/kras:g12d**) and KRAS:G12C (**/kras:g12c**). Alternatively, mutations can be aggregated at the residue level so that a combined ***switch*** analysis is performed for all KRAS:G12 mutations (**/kras:g12?**), or at the gene level so that a combined ***switch*** analysis is performed for all KRAS GoF mutations in the curated dataset (profile **/kras:a146p,a146t,a146v,a59t,g12a,g12c,g12d,g12r,g12s,g12v,g13c,g13d,k117n,q61h,q61k,q61l,q61r**). While hits obtained with aggregated GoF profiles may need further analysis to tease out specific contributions of individual mutations, they are an effective way of increasing the number of observations so that sufficient (>5) are obtained for the significance calculation. A full ***switch*** screen of our curated GoF dataset using gene-level aggregation identified 2082 significant hits with strong SSL to a GoF mutated gene, and took under 2 hours. The full hit-list as well as the aggregated profile definitions used is provided in **Supplementary Database 2**.

Some of the strongest signals in the ***GOF*** analysis occur between GoF mutated genes and themselves (**Figure 12**). Most of the genes that display this effect are well known highly penetrant oncogenes e.g. KRAS, BRAF, NRAS, PIK3CA, CTNNB1, NOTCH1, ALK – and the self-dependence reflects the phenomenon of oncogene addiction (Weinstein & Joe, 2008) that makes the products of these genes compelling targets for drug discovery (Torti & Trusolino, 2011). Many of the genes identified by non-self signals in the ***GOF*** analysis are transcription factors mediating the downstream effects of the activated oncogene with which they consequently display an SSL relationship, but these tend to be very difficult targets for drug discovery.

One notable exception is CRAF (RAF1), a druggable target showing a significantly SSL effect with activated NRAS, and weaker but nonetheless significant SSL effects with KRAS and HRAS (**Figure 13**). Given the challenge in directly targeting NRAS, CRAF inhibitors (of which several are available) would be expected to be useful in NRAS activated tumours (Bekele *et al*, 2021). Despite the significant degree of functional redundancy between CRAF and BRAF, the effect is restricted to CRAF, with BRAF showing negligible SSL with any of the RAS genes.

**Figure 5.**
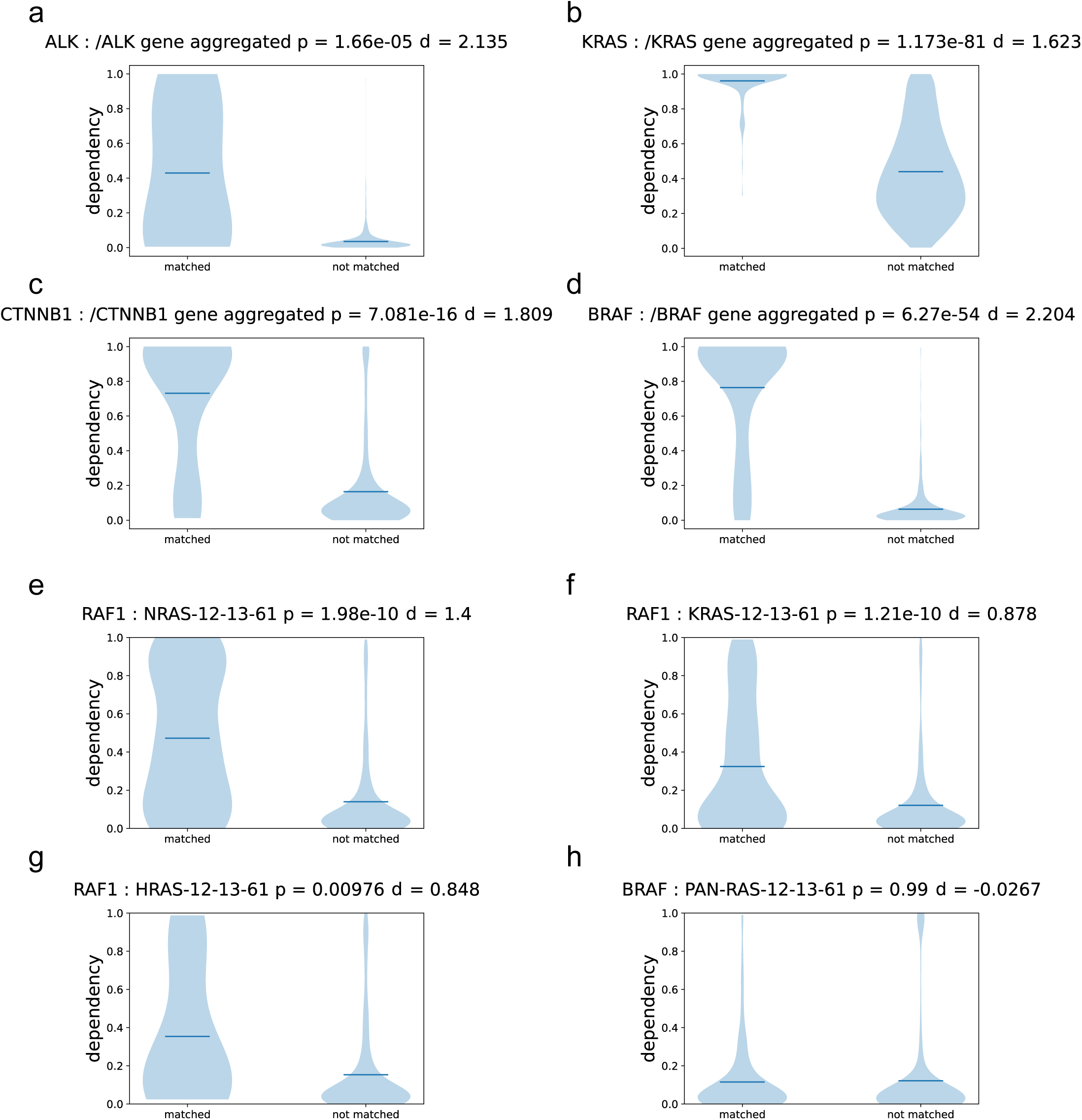
**a)** KDE plot of strong self-dependencies found in the ***GOF*** analysis demonstrating cellular ‘addiction’ to ALK. Definitions of the gene-aggregated GoF profiles used are listed in **Supplementary Table 1**. **b)** as **a)** but for KRAS. **c)** as **a)** but for CTNNB1 **d)** as **a)** but for BRAF. **e)** KDE plot of CRAF (RAF1) dependency in cell lines harbouring activating mutations at residues 12,13 or 61 in NRAS. **f)** as **e)** but for KRAS. **g)** as **e)** but for HRAS. **h)** KDE plot of BRAF dependency in cell lines harbouring activating mutations in any RAS paralogue. Unlike CRAF (RAF1), BRAF shows no SSL with mutations in any of the RAS genes.

### Mining for Secondary Factors

While strong effects of mutational disruption/deletion of individual genes on the dependency of other genes can be detected by pairwise analyses (Lord *et al*, 2020), these are relatively unusual, and in most circumstances multiple factors play a role in determining whether a given gene is important or not in the survival of a cancer cell line (Ryan *et al*, 2023a). Expanding pairwise relationships to a third dimension is feasible but the computation required increases very substantially, and the overwhelming majority of the triple combinations examined will not be informative. The ability of *DepMine* to define cancer profiles allows it to leverage the significant body of prior knowledge available in cancer biology and identify genes whose dependency switches as a result of far more complex situations than the loss of function of a single gene.

For example, the **APC_LOSS** profile that generates a strong and highly significant SSL with CTNNB1 (β-catenin) (**Figure 2d**), nonetheless matches a number of cell lines that display a low dependency. Applying the *DepMine* mining algorithm to this data we find that profile-matching cell lines deriving from paediatric samples or of uterine or lymphoid origin are significantly over-represented in this low-dependency population (**Figure 6a,b,c**), while those originating from bowel are significantly over-represented in the high-dependency population (**Figure 6d**). Given the high prevalence of mutations, copy number and expression level changes in cancer cell lines in general, mining for secondary genes is restricted to changes that occur in more than a specified fraction of the cell lines matching the profile – 10% empirically provides a useful cut-off. A significant strong effect was observed for GoF mutation of KRAS (**Figure 6e**) which occurs in nearly 1/3 of the matching cell lines and is associated with high CTNNB1 dependence, and a significant strong effect was found for deletion of CDKN2A which is associated with low dependency on CTNNB1 (**Figure 6f**).

**Figure 6.**
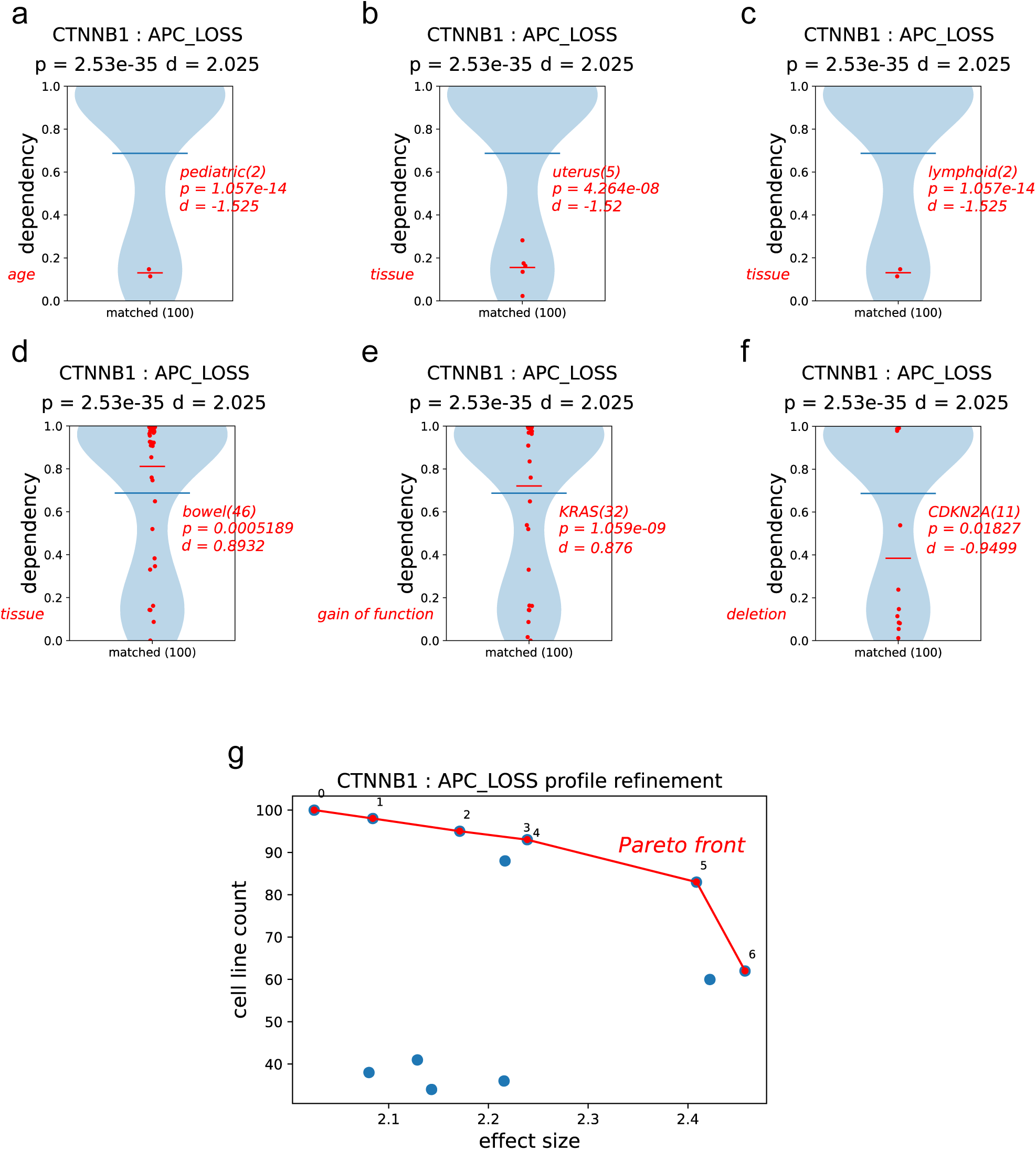
**a)** KDE plot for dependency on CTNNB1 in cells matching the **APC_LOSS** profile, overlayed with distribuRons of cell lines of paediatric origin displaying significant enrichment in the low dependency sub-populaRon (red). **b)** as **a)** but for cell lines derived from uterus, displaying significant enrichment in the low dependency sub-populaRon (red). **c)** as **a)** but for cell lines derived from lymphoid Rssues, displaying significant enrichment in the low dependency sub-populaRon (red). **d)** as **a)** but for cell lines derived from bowel, displaying significant enrichment in the high dependency sub-populaRon (red). **e)** as **a)** but for cell lines KRAS GoF mutaRons, displaying significant enrichment in the high dependency sub-populaRon (red). **f)** as **a)** but for cell lines with deleRon of the CDKN2A gene, displaying significant enrichment in the low dependency sub-populaRon (red). **g)** Plot of cell line count versus effect size for dependency on CTNNB1 gene by cells selected by the APC_LOSS profile with addiRon of different combinaRons of secondary factors. The red line and numbering indicate Pareto opRmal soluRons.

### Automatic generation of complex profiles

Application of the Pareto optimisation function (Hu *et al*., 2013) in *DepMine* analysed all possible combinations of these secondary factors that were individually found to influence the CTNNB1 dependency profile in cells with APC^LoF^, and identified optimal combinations with respect to effect size and the number of cell lines they match (**Figure 6g**). Combination 5 (**CTNNB1_APC_LOSS_REF_5**), which combines APC^LoF^ with tissue specifications that exclude cells of paediatric origin or from uterine or lymphoid tissue, and excludes cells in which CDKN2A (which encodes p16^INK4A^ and p14^ARF^) is deleted, achieves a substantial increase in effect size, while retaining applicability to 83 cell lines and probably constitutes a globally optimal solution. Combination 6 (**CTNNB1_APC_LOSS_REF_6**) which doesn’t apply the restriction on CDKN2A but adds a positive requirement for a bowel tissue origin in its place, further improves the effect size, but decreases the cell line cover to only 62. The final choice of cell line selection in which to explore inhibition or targeted depletion of CTNNB1, is up to the investigator and will depend on whether they are more concerned with the breadth of cancers that can be targeted or the size of the effect likely to be elicited.

In a second example we took the observation of a strong SSL relationship between WRN and our microsatellite instability profile **MSI_FULL_DOWN** (**Figure 7a**) and mined the profile matching cell line population for secondary factors. The ***mine*** algorithm identified two tissue-origins*: lymphoid*, strongly associated with low WRN dependency, and *bowel*, strongly associated with high WRN dependency, which effectively subdivide the profile-matching cell population (**Figure 7b,c**). Indeed, this is essentially the selection criteria for ongoing clinical trials of WRN inhibitors (Moschetta *et al*, 2023) - however it is not identified as an optimal profile in our analysis. Six individual genes were also identified whose mutational disruption in > 20% of the cell lines matching the **MSI_FULL_DOWN** profile showed significant strong effects associated with high WRN dependency : ARID1A a component of SWI/SNF chromatin remodelling complexes mutated in >5% of cancers (Mullen *et al*, 2021), KMT2B and KMT2D are lysine methyltransferases frequently mutated in a range of cancer (Rao & Dou, 2015). Of lesser impact are RPL22 - a ribosomal protein whose loss has been associated with oncogenesis through multiple mechanisms (Goudarzi & Lindstrom, 2016); PTEN - a lipid phosphatase tumour suppressor that negatively regulates signalling through the PI3K/AKT axis (Lee *et al*, 2018); and TTN, which encodes titin – the largest protein in the human genome with ∼34,000 amino acids. Although TTN is not directly linked to any tumour suppressor pathway, mutation within the TTN gene has been shown to be an effective indicator for overall mutational burden in a tumour cell line and indicative of MSI in colorectal cancer samples (Oh *et al*, 2020).

**Figure 7.**
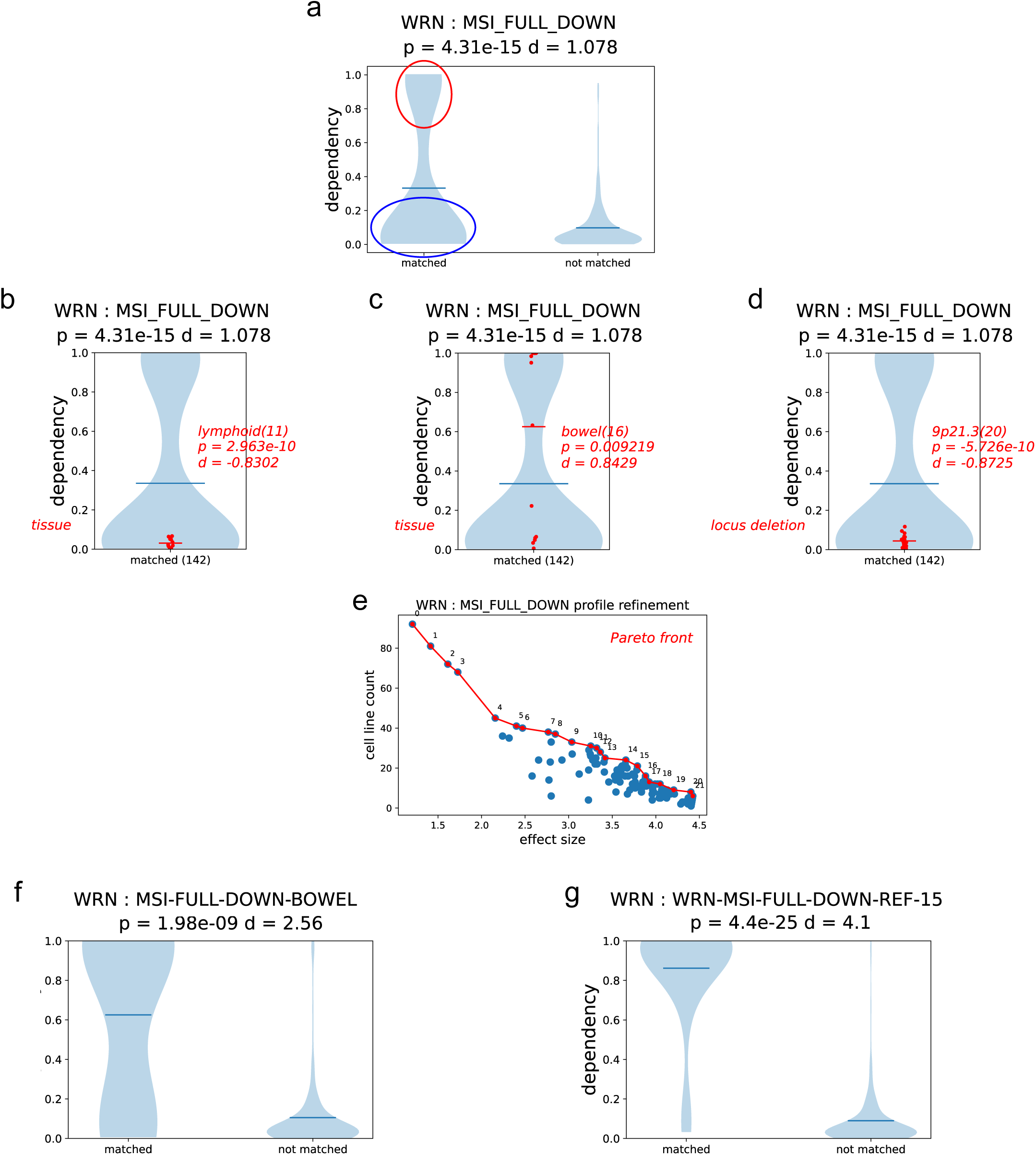
**a)** Within the distribuRon of cell lines matching the **MSI_FULL_DOWN** profile, there are disRnct sub-populaRons which show high dependency on WRN (red oval) and those that show low dependency (blue oval). **b)** KDE plot for dependency on WRN in cells matching the **MSI_FULL_DOWN** profile, overlayed with distribuRons of cell lines displaying significant enrichment in the low dependency sub-populaRon (red) of cells derived lymphoid Rssue. **c)** as **b)** but for cell lines displaying significant enrichment in the high dependency sub-populaRon (red) of cells derived from bowel **d)** as **b)** but for cell lines displaying significant enrichment in the high dependency sub-populaRon (red) of cells with deleRon of cytogeneRc locus 9p21.3. **e)** Plot of cell line count versus relaRve effect size for dependency on WRN in cells selected by the **MSI_FULL_DOWN** profile with addiRon of different combinaRons of secondary factors. The red line and numbering indicate Pareto opRmal soluRons. **f)** KDE plot of WRN dependency in cell lines matching the **MSI_FULL_DOWN** profile, with the addiRonal specificaRon of a bowel Rssue origin, compared with the control cell populaRon. **g)** as **f)** but with cells selected by profile combinaRon 15 from the automaRc opRmisaRon process.

Mining also identified two copy number variations : deletion of a region of 9p21.3 containing CDKN2A, CDKN2B and MTAP is strongly associated with low dependency on WRN (**Figure 7d**); and amplification of a region of 4p16.1 containing the uncharacterised FAM90A26, DEFB131A encoding a β-defensin isoform, and an extremely unusual multi-gene cluster encoding 20 members of the USPL17 family of deubiquitinating enzymes (Burrows *et al*, 2005), was associated with high WRN dependency.

In contrast to the CTNNB1-APC^LoF^ situation where some of the factors are redundant and several combinations have similar effect size and cell count scores, the combinatorial exploration of the 1024 decision trees built from the 10 secondary factors identified for the **MSI_FULL_DOWN** profile give a complex distribution (**Figure 7e**). These are divided into a broad-acting subgroup with small improvements of the original profile, and a large poorly differentiated cluster with substantially higher effect size, but selecting very much smaller cell line populations. The effectiveness of the ***mine*** algorithm can be clearly seen by comparing the switching analysis for a ‘hand-made’ profile **MSI_FULL_DOWN_BOWEL**, where the clinical trial recruitment requirement for a bowel tissue origin is added onto the base MSI profile (**Figure 7f**), with one of the Pareto refined profiles from ***mine*** (solution 15 in **Figure 7e**) that additionally specifies avoidance of tumours of lymphoid origin or with 9p21.3 deletion, and has positive requirements for LoF mutations of KMT2D and TTN on top of the basic MSI profile (**Figure 7g**). Not only does the refined solution deliver a substantially improved effect size, but it matches 21 DepMap cell lines with dependency data compared to 16 for the tissue focussed profile.

## DISCUSSION

Cancer gene dependency screens, exemplified by DepMap (Tsherniak *et al*., 2017) are rich sources of information with the potential to accelerate identification of new therapeutic targets, and at least as important, identify selective genetic and epigenetic biomarkers that enable targeted therapies to be used with maximal effect. However, the complexity of such data and its associated metadata, especially in their raw state, limits the depth and breadth to which the average laboratory researcher can explore, and typically this is restricted to browsing data on ‘favourite’ genes. Analysis that takes on the entirety of the data and metadata requires substantial computational and data-science expertise, and while of considerable value, these studies do not always ask the questions the laboratory researcher or clinician might wish to ask. DepMap itself provides some powerful query and analysis tools on its website https://depmap.org/portal/, and other groups have developed independent interfaces and packages that provide additional or complementary analysis (Killian & Gatto, 2021; Shimada *et al*, 2021; Wappett *et al*, 2021).

The *DepMine* software presented here seeks to provide a comprehensible but still very powerful toolkit with which highly complex questions can be posed across the entire cell and gene dependency collection, without the need for computational/data science expertise beyond the ability to frame a *cancer profile* as a basic algebraic formula. The breadth of criteria interrogated by the primitive postulate operators allows selection on almost every useful characteristic of a cancer cell line, and these can be elaborated into *cancer profiles* of essentially unlimited complexity using basic set operators and brackets. Defined *cancer profiles* can be assembled into collections, allowing the compilation of a library of cell line definitions that can be used for multiple downstream analyses. Although based on the data architecture of DepMap, the *DepMine cancer profile* selection system is fully adaptable to any cell line collection – public or private – where mutational, copy-number, and expression data has been generated.

## ACKNOWLEDGEMENTS

We are grateful to our colleagues for critical discussions during the development of *DepMine*, and for presenting us with challenging problems to solve, which have helped drive its development. This work was supported by The University of Sussex and The Institute of Cancer Research.

## AVAILABILITY

The Python implementation of *DepMine* and associated data files can be obtained by request from and is free to academics and Not-For-Profit organisations.

## Supplementary Data

### Data Files

DepMine uses a number of datafiles which are either supplied with the code or can be downloaded from a range of different sources :

| File | Data | Web Page | Version |
| --- | --- | --- | --- |
| CRISPRGeneDependency.csv | gene dependency | https://depmap.org/portal/download/all/ | Public 23Q2 |
| Model.csv | cell line information | https://depmap.org/portal/download/all/ | Public 23Q2 |
| OmicsSomaticMutations.csv | mutations | https://depmap.org/portal/download/all/ | Public 23Q2 |
| OmicsCNGene.csv | copy number | https://depmap.org/portal/download/all/ | Public 23Q2 |
| OmicsFusionFiltered.csv | gene fusions | https://depmap.org/portal/download/all/ | Public 23Q2 |
| OmicsExpressionProteinCodingGenesTPMLogp1.csv | expression | https://depmap.org/portal/download/all/ | Public 23Q2 |
| c1.all.v2023.1.Hs.json.txt | chromosomes | https://data.broadinstitute.org/gsea-msigdb/msigdb/release/2023.1.Hs/ | 2023.1.Hs |
| hgnc_complete_set.json | HUGO gene names | https://www.genenames.org/download/archive/ | Current |
| c2.cp.reactome.v2023.2.Hs.symbols.gmt | pathways | https://data.broadinstitute.org/gsea-msigdb/msigdb/release/2023.2.Hs/ | 2023.2.Hs |
| Homo_sapiens.GRCh38.110.refseq.tsv | ENSEMBLE reference sequences | https://www.ensembl.org/Homo_sapiens/Info/Index | GRCh38 |
| ChromosomeLengths.csv | Chromosome lengths | local |  |
| ExpressionThresholds.csv | Expression thresholds | local |  |
| Assignment_Solution_Activities.tsv | Mutation signature assignments | local |  |
| Final_GOF_all_table | Annotated GoFs | local |  |
| AB_Dep_Profiles | cancer profiles | local |  |

**Supplementary Table 1 -.**
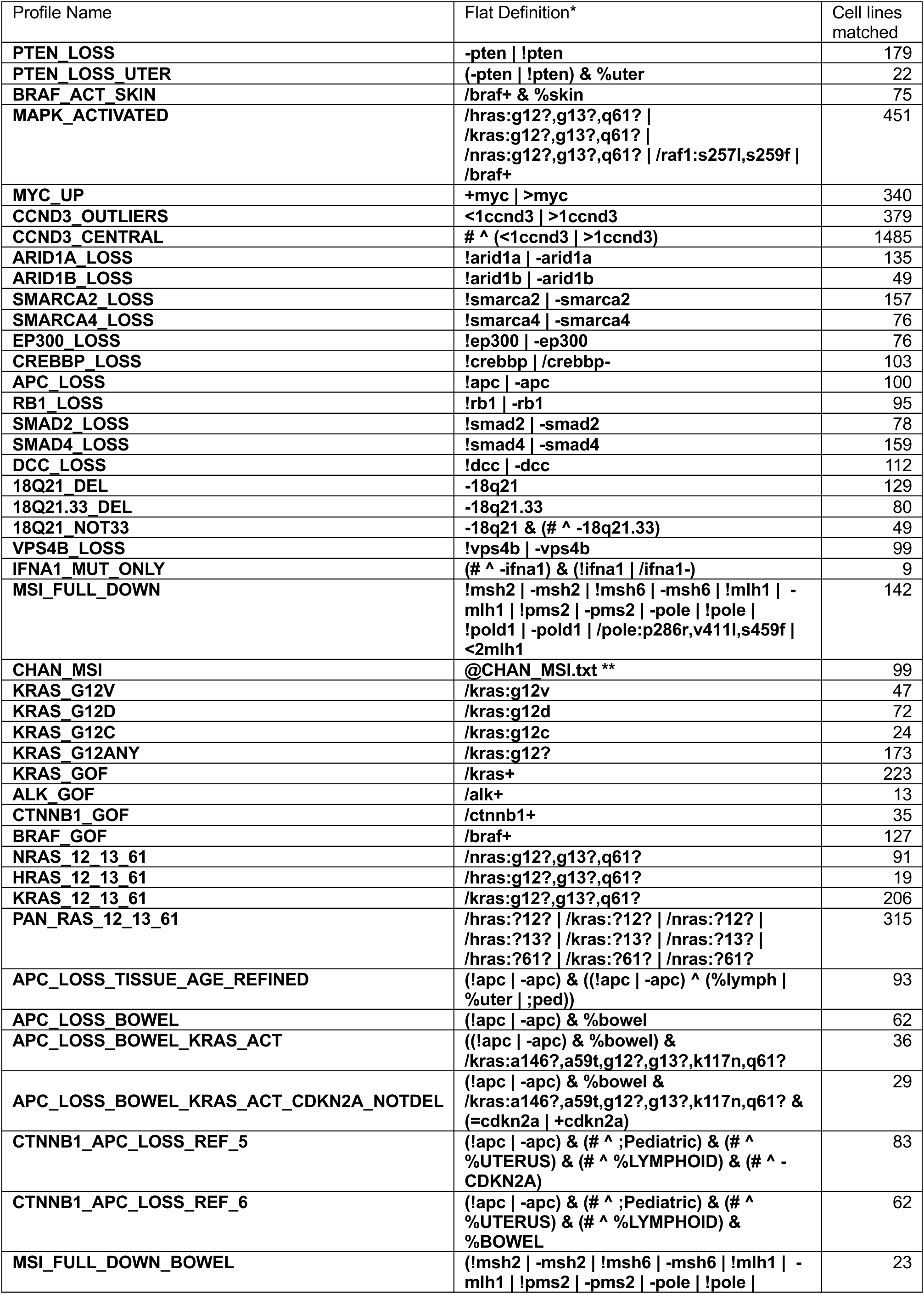
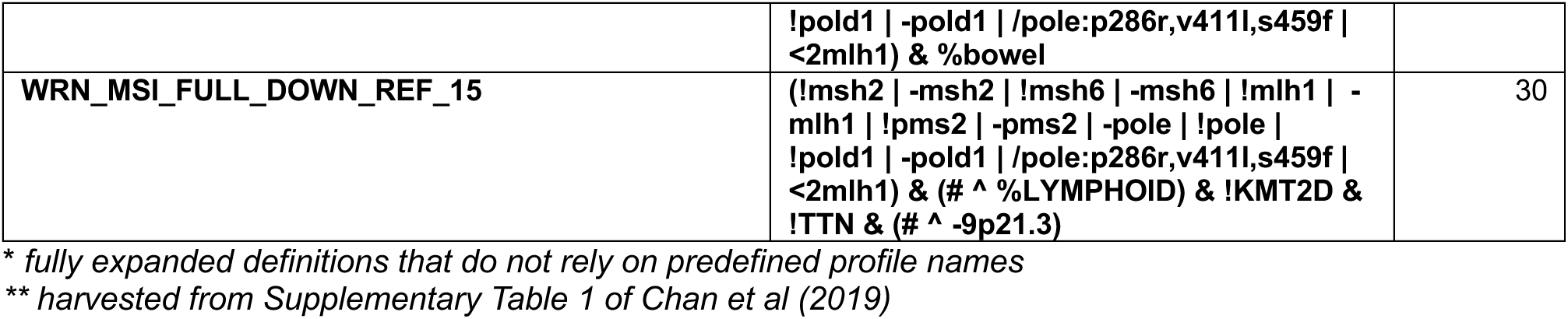
Table of cancer profile definitions referenced in the main text.

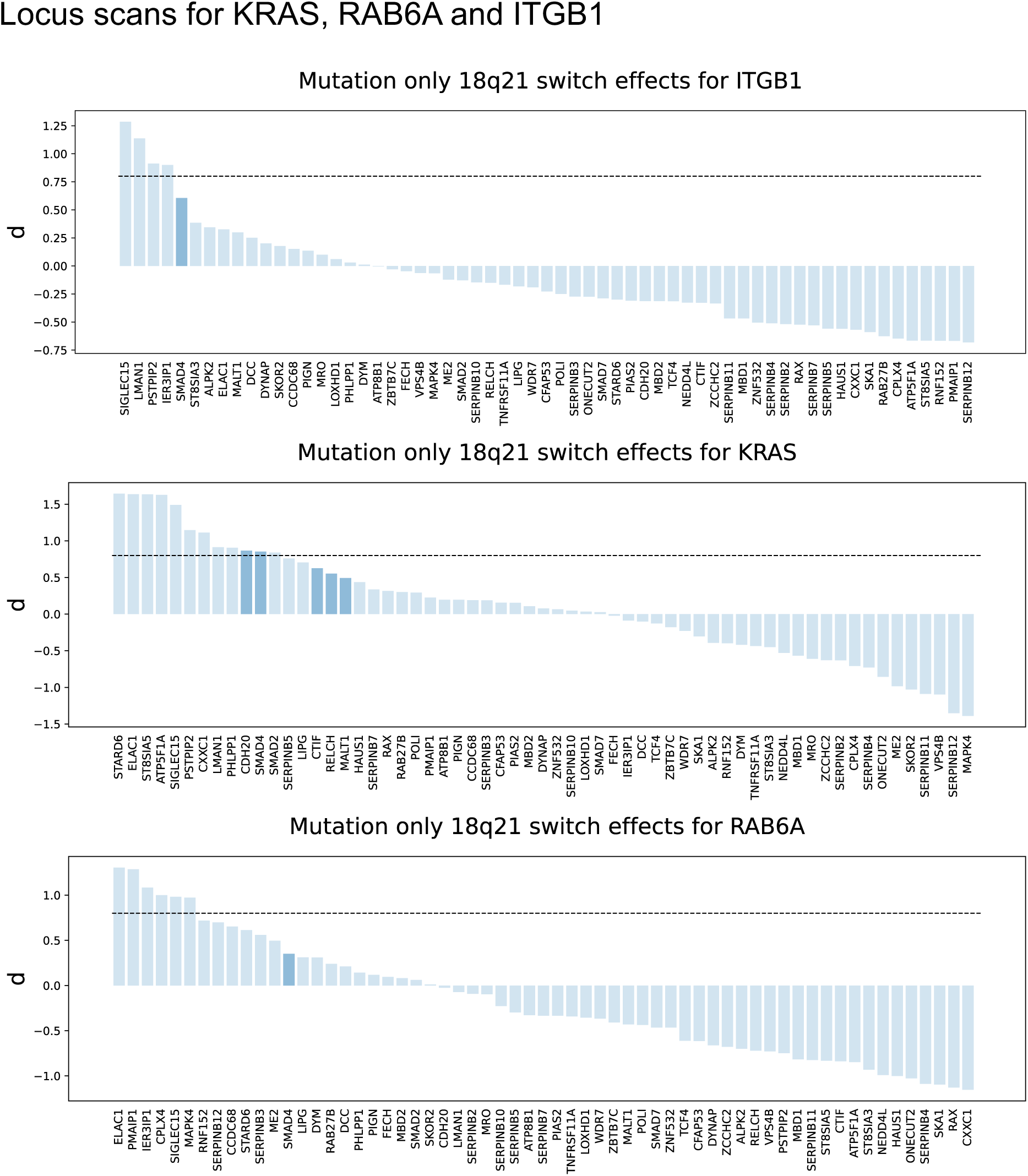

## Supplementary Databases

**Supplementary Database 1.tsv** – List of 34256 GeneA:GeneB pairs where the |dep(GeneA)| < 0.65, and dep(GeneA) in cells with LoF/CNV-del of GeneB was significantly > than GeneB-WT cells (p < 0.05).

**Supplementary Database 2.tsv –** List of 2082 GeneA:GeneB pairs where GeneB has any literature annotated Gain-of-Function (GoF) mutation, and where dep(GeneA) in cells with GeneB-GoFwas significantly > than GeneB-WT cells ((p < 0.05 and d > 0.8).

